# Neural pathways linking hypoxia with pectoral fin movements in *Danio rerio*

**DOI:** 10.1101/655084

**Authors:** Kaila Rosales, Chen-Min Yeh, Javier J. How, Reginno Villa-Real, Elizabeth DePasquale, Alex Groisman, Sreekanth H. Chalasani

**Author notes:** Author for correspondence (S.H.C.). These authors contributed equally to this work (Kaila Rosales, Chen-Min Yeh).

## Abstract

Zebrafish larvae respond to hypoxia by increasing a number of ventilatory behaviors. During development, these animals switch from skin-resident to gill-resident neuroendocrine cells around 7 days post fertilization (d.p.f.) to detect hypoxia and drive adaptive behaviors. Here, we probe the neural pathways that receive inputs from skin-resident neuroendocrine cells and alter pectoral fin movements. We first show that a 5 d.p.f. larva increases its pectoral fin movements and heart activity upon hypoxia exposure. Next, we map the downstream neural circuitry and show that individual vagal sensory neurons receive inputs from multiple oxygen-sensing neuroendocrine cells. We then use calcium imaging to show that neurons in the second, but not third, vagal sensory ganglia show increases in the magnitude of their hypoxia-evoked responses. Finally, we link purinergic signaling between neuroendocrine cells and second vagal sensory neurons to increases in pectoral fin movements. Collectively, we suggest that vagal sensory neurons transform hypoxic stimuli into respiratory behaviors.

## Introduction

Animals sense changes in their ambient oxygen levels and alter their behaviors to maintain their internal oxygen concentration within a normal physiological range. In mammals, the type I cells of the carotid body and neuroepithelial bodies in the lungs detect changes in environmental or arterial O_2_ and CO_2_ concentrations and initiate compensatory changes in ventilation and heart rate [1-3]. In teleost fish, the O_2_- and CO_2_- sensitive neuroendocrine cells (NECs) of the gills are homologues of these mammalian chemoreceptors. Zebrafish gill NECs respond to acute hypoxia and high P_CO2_ [4, 5] and contain synaptic vesicles that are believed to be released upon stimulation onto afferent fibers of the glossopharyngeal and vagus nerves [6, 7]. In addition, during development these animals transition from anoxia-tolerant embryos to hypoxia-sensitive larvae within 2-3 days post-fertilization (d.p.f.) [8], but lack innervated gill NECs until 7 d.p.f. [9]. Despite this, 3-7 d.p.f. larvae show robust hyperventilatory responses to hypoxia [9], which are mediated by a population of skin NECs [10]. However, it is not clear whether the skin and gill NECs share common downstream neural circuitry.

During development, the underlying neural circuits affect different behavioral adaptations to hypoxia. For example, the anoxia-tolerant 2 d.p.f. embryos can alter their frequency of pectoral fin and body movements after exposure to hypoxic solution. After hatching, 3 d.p.f. larvae show a significant increase in their rate of buccal and opercular movements (collectively termed ventilation behavior) during hypoxia. This response is irregular in frequency, but was found to be synchronous with the movement of pectoral fins [9, 10]. In addition, a role for pectoral fin movements in cutaneous gas exchange was also identified using ablation studies [11, 12]. While adult and juvenile pectoral fin swimmers increase their fin beat frequency to achieve greater swim speeds [13, 14], zebrafish larvae show no change in pectoral fin beats during rapid swimming [12]. Instead, pectoral fin movements were found to bring distant fluid toward the body and move it caudally behind the fins disrupting the boundary layer along the animal’s skin, a site for oxygen absorption in larvae [12]. Consistently, we and others have shown that 5 d.p.f larvae increase their pectoral fin movements upon hypoxia exposure [12, 15-17].

While gill NECs synapse onto afferent fibers of the glossopharyngeal and vagal nerves likely using catecholamine and serotonin neurotransmitters [6, 18], little is known about the signaling between the skin NECs and their downstream neural circuitry. This is particularly relevant to our understanding of how the hypoxia sensing-neural circuits are altered during development, matching the demand of the growing animal. We use sparse labeling in transgenic larvae to show that the skin NECs are innervated by vagal sensory neurons. Next, we combine calcium imaging, pharmacology and cell ablations to reveal a role for purinergic signaling between skin NECs and neurons of the second vagal ganglion in modifying pectoral fin movements. Together, this work reveals a potential circuit involved in ventilatory behaviors in larval zebrafish

## Results

### Larval zebrafish increase their pectoral fin movements and heart activity upon hypoxia

Previous studies have identified a role for pectoral fin movements in mediating larval responses to hypoxia [12, 16]. Additionally, zebrafish larvae have also been shown to increase their heart rate in response to hypoxia [19]. We previously used a microfluidic device that rapidly reduced the level of oxygen in the media around the larva [15]. In that device, the larvae were relatively free to move, making it poorly suited for obtaining imaging data at cellular resolution, which would be required for probing the underlying neuronal circuits. To overcome this challenge, we designed a new microfluidic-based device where the trunk and the head of a 5 d.p.f. larva are trapped in an agarose gel (Figure 1A, 1B). This device allows us to monitor both pectoral fin and heart movements (Figure 1C). We observed that larvae increase their rate of pectoral fin movements to both strong and weak hypoxia and heart activity to strong hypoxia alone (Figure 1C-1E, Supplementary Videos S1, S2). To test whether these behavioral changes are specific to hypoxia we also analyzed larval responses to a different stressor, osmotic shock. We found that larvae exposed to water containing high amounts of sodium chloride, which is known to increase cortisol levels in larvae [20], which reduces pectoral fin movements and increases heart activity (Figure 1F, 1G). Taken together, these data suggest that the increased rates of pectoral fin movements are specific to hypoxia-induced stress.

**Figure 1.**
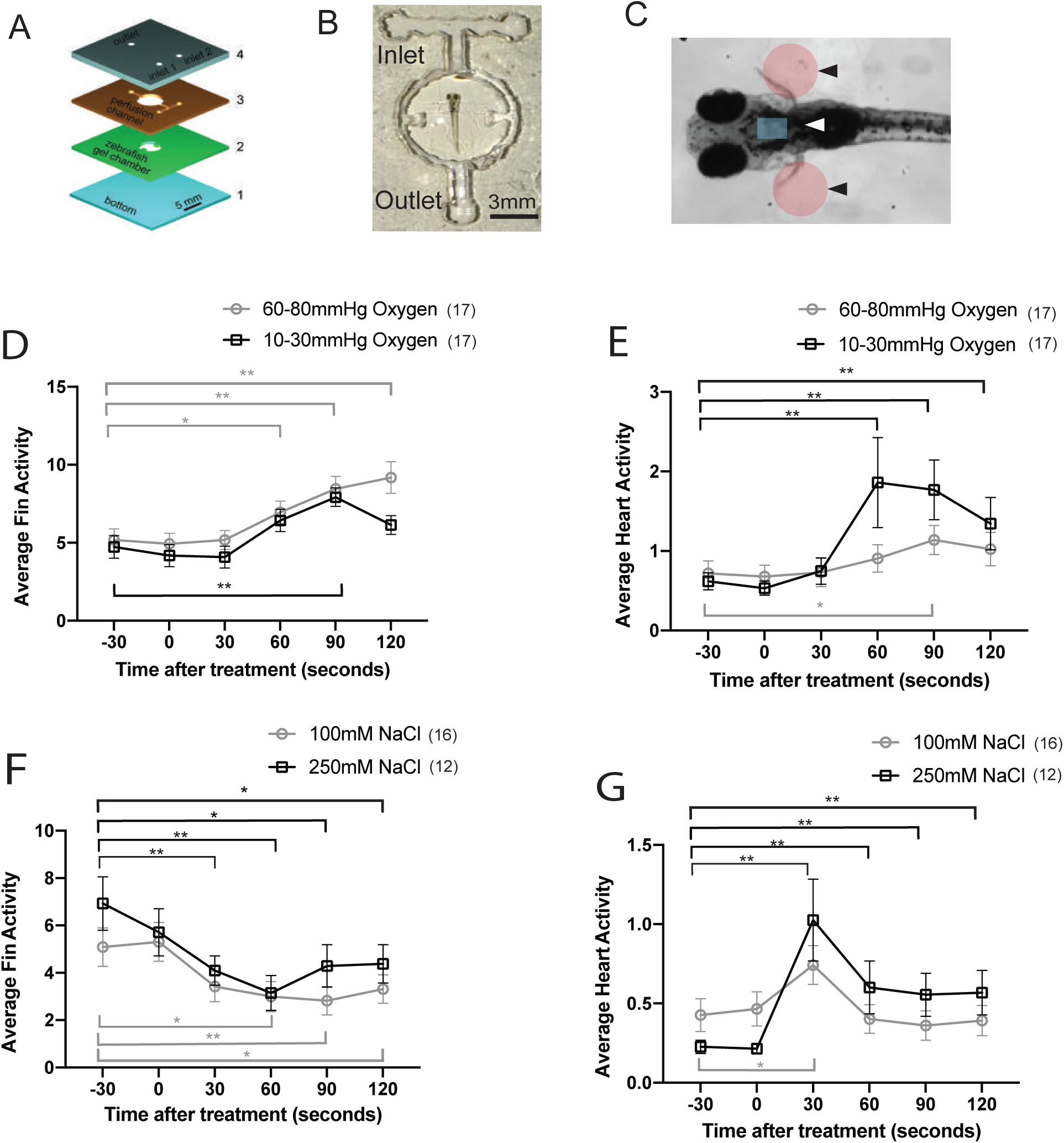
Hypoxia induces respiratory behaviors in zebrafish larvae. **A**., Schematic showing the laminated microfluidic device made of acrylic layers with a coverglass bottom. **B**., Photograph of the microfluidic device with a zebrafish larva embedded in agarose gel in the chamber. **C**., Zebafish larva with shaded circles (black arrowheads) and rectangles (white arrowheads) identifying pectoral fin and heart movements, respectively. Average (**D**.,) fin activity and (**E**.,) heart activity in zebrafish larvae exposed to hypoxia levels of 10-30 and 60-80 mmHg at t =0. Normoxia is 150 mmHg. Zebrafish larvae were exposed to different concentrations of NaCl (osmotic shock) at t = 0 and their (**F**.,) fin and (**G**.,) heart activity was measured. Averages and s.e.m. are shown in (**E-H**), n > 12 with *p< 0.1 and ** p< 0.05 using one sample t-test.

### Skin NECs send afferents onto neurons of cranial sensory ganglia

In the 5 d.p.f. larvae, the skin NECs can be identified by serotonin immunolabeling and are distributed over the entire skin [10]. We hypothesized that these skin NECs would synapse onto neuronal afferents from one or more of the four cranial sensory ganglia (trigeminal, facial, glossopharyngeal, and vagal) that are known to transduce somatosensory, chemosensory and viscerosensory information from receptors in head, throat, heart and body viscera to the brain [21]. To test our hypothesis, we used transgenic animals where GFP expression was driven by the promoter of a P2X(3) purinoceptor subunit and labels neuronal cell bodies and processes of these cranial ganglia [22]. We were able to identify the cell bodies of the glossopharyngeal ganglion (IX), and the three sub clusters of the vagal ganglia (X-1,X-2, and X-3) (Figure 2A, 2B). We double-labeled tg(*p2xr3*.*2::eGFP*^*sl1*^) with anti-*gfp* and anti-serotonin antibodies and found that skin NECs appear to be innervated by dendritic processes from neurons in both glossopharyngeal and vagal ganglia (Figure 2C). These results suggest that skin NECs, particularly those that are close to the gills in the 5 d.p.f. larvae are innervated by glossopharyngeal and vagal neurons.

**Figure 2.**
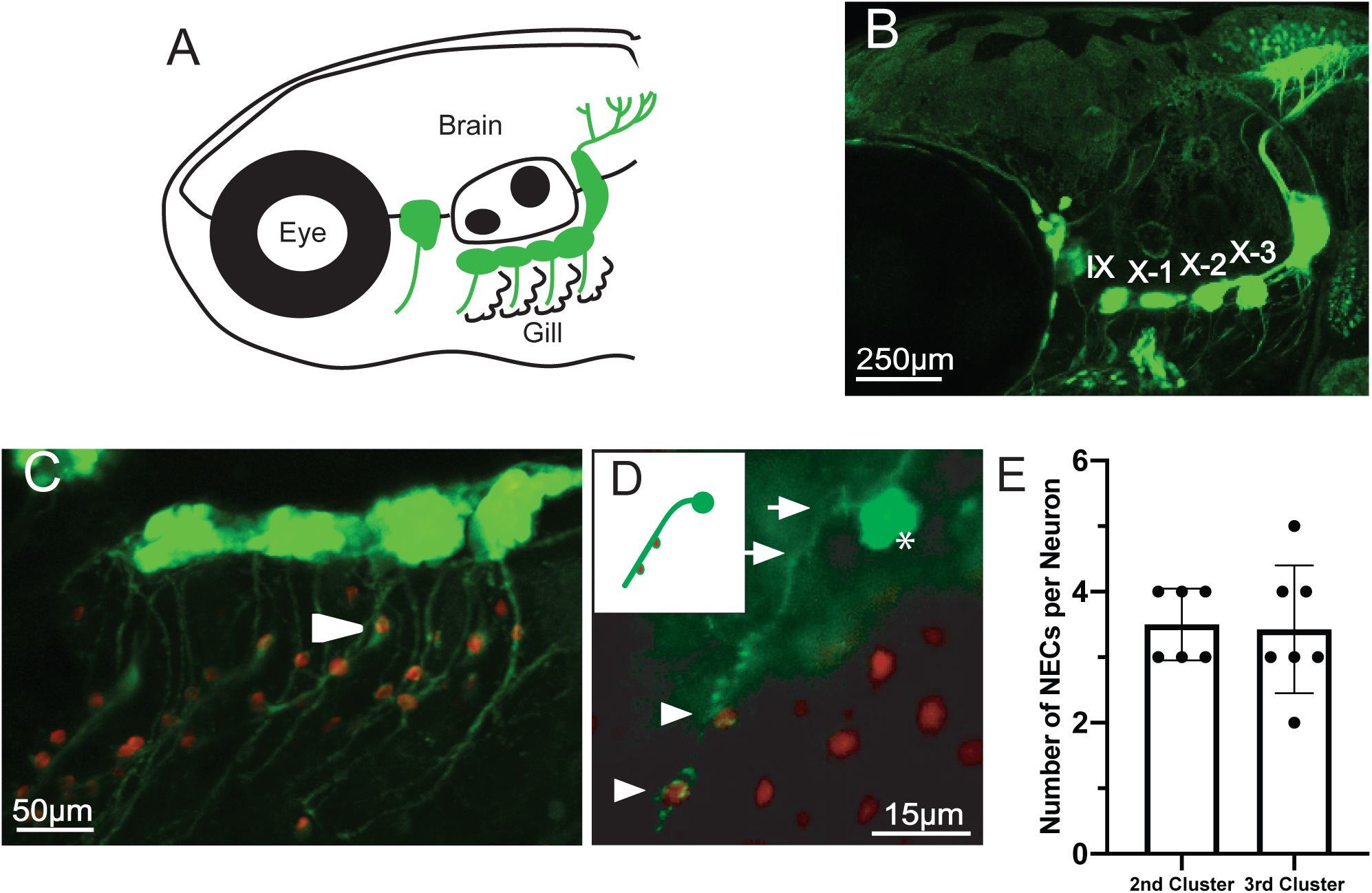
A many-to-one circuit encodes oxygen levels. **A**., Schematic showing the lateral view of a zebrafish larva with cranial sensory neurons and their projections labeled in green. **B**., Lateral view of a *Tg(p2rx3*.*2:gfp)* transgenic larva with neurons in the IX and X cranial sensory ganglia expressing GFP. **C**., Immunohistochemistry showing a Tg(p2rx3.2:gfp) transgenic larva with cranial sensory neurons (green) and neuroendocrine cells (red, NECs) labelled using anti-gfp (green) and anti-serotonin (red) antibodies respectively. Arrowhead indicates an example of cranial nerve afferent wrapping around NECs. **D**., Immunohistochemistry showing the afferent nerve of an individual vagal sensory neuron (green) wrapping around multiple neuroendocrine cells (red) in a *PB4; UAS:Kaede* transgenic larva labeled using anti-kaede (green) and anti-serotonin (red) antibodies. Asterisk, arrowheads, and arrow indicate the neuron, contact sites between NECs and vagal afferents, and neuronal projection respectively. Inset is the schematic of the neuron. **E**., Number of NECs wrapped by a single afferent nerve of neurons in either the second or third vagal cluster. Each dot represents an individual vagal sensory neuron; Columns are mean value; bars are S.D.

Next, we tested the relationship between skin NECs and vagal sensory neurons using sparse labeling. We crossed a tg(*PB4:GAL4*) fish with tg(*UAS::kaede*) fish to obtain embryos where the fluorescent protein kaede is expressed in populations of cranial sensory neurons [23]. We observed a range of kaede expression in the progeny of this cross with some embryos showing labeling in only a few cranial sensory neurons, while others labeled the entire population. We sorted this population to identify those embryos where *kaede* expression was limited to one or a few vagal sensory neurons and labeled these animals with anti-kaede and anti-serotonin antibodies. We observed that individual vagal sensory neurons receive inputs from multiple NECs (Figure 2D). We also quantified the number of vagal sensory neurons and contacting NECs across multiple animals and found that individual vagal sensory neurons received inputs from 2-5 skin NECs (Figure 2E). Taken together, these studies suggest that multiple skin NECs send inputs to vagal sensory neurons in the 5 d.p.f. larva.

### Second, but not third, vagal ganglionic neurons respond to hypoxia

To study how the downstream vagal sensory ganglia responded to hypoxia, we used a light sheet microscope to record calcium responses in the neurons of both the 2^nd^ and 3^rd^ vagal sensory ganglia in larvae expressing neuronal nuclei-localized GCaMP6f [24] (it is technically difficult to record from the 1^st^ vagal ganglia). We observed recurring calcium events lasting a few seconds that were particularly prevalent in neurons of the second compared to third vagal sensory ganglion (Figure 3C, Supplementary Figure S1, S2, Supplementary Videos S3, S4).

**Figure 3.**
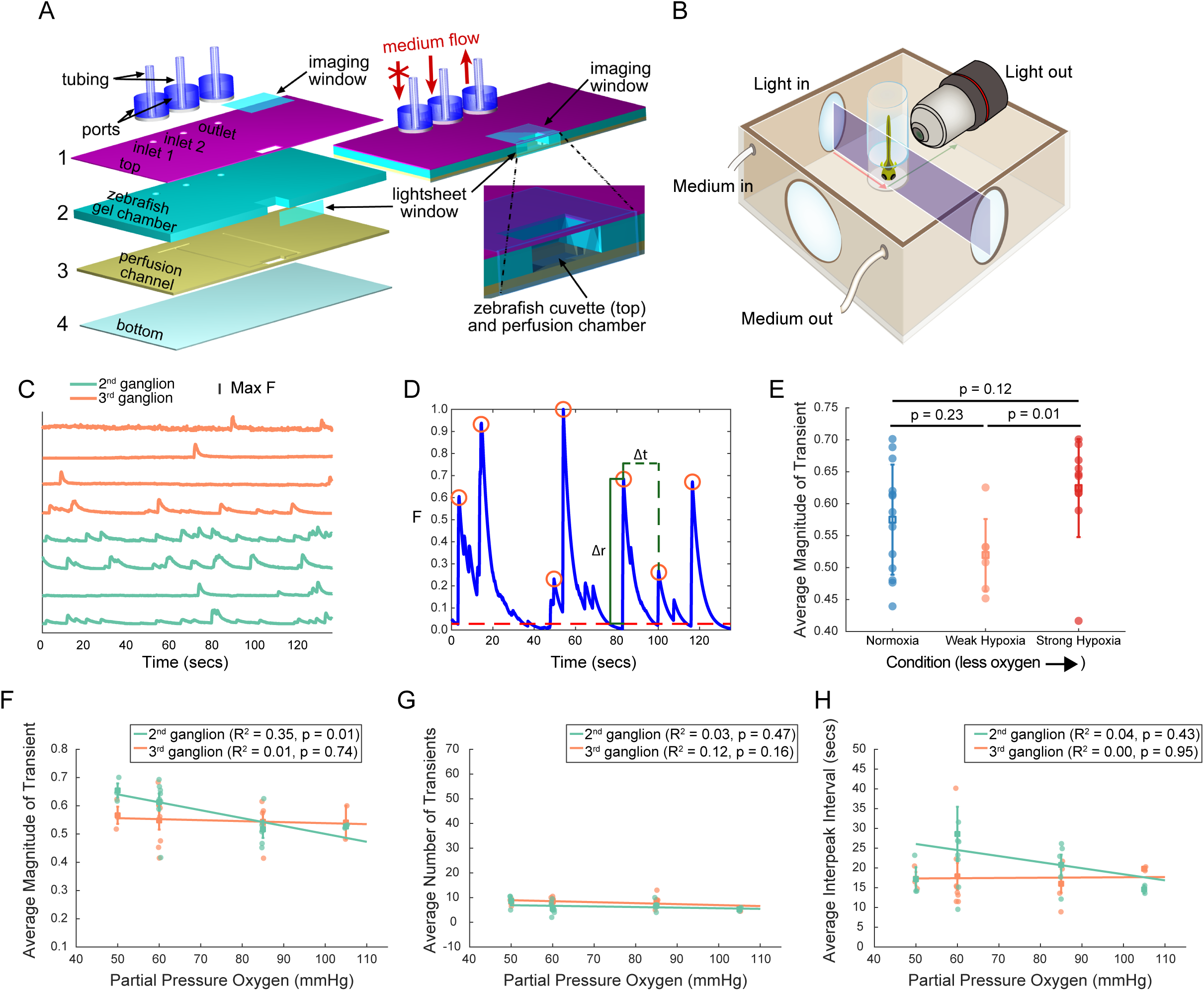
Calcium imaging reveals correlation between hypoxia stimuli and magnitude of neural responses in the 2nd, but not 3rd vagal sensory ganglion. **A**., Schematic of the microfluidic device with rapid medium exchange for light-sheet microscopy imaging of a zebrafish larva to detect neural activity under acute hypoxia. **B**., Illustration of light-sheet microscopy. **C**., Example traces, normalized to peak fluorescence, from ganglion 2 (green) and 3 (orange) of one larva. **D**., Detecting and validating peaks from the GCaMP time series of a neuron. The red circles mark peaks, the red dashed line is the baseline (10th percentile of normalized fluorescence, F), the dashed green line (Δt) indicates the time between peaks (i.e. interpeak interval) and the solid green line (Δr) indicates the value of the peak (i.e. magnitude). The trace is normalized by its largest attained magnitude. In the example, and throughout our analysis, we used a weight of 0.7 to validate peaks. **E**., The average magnitude of calcium transients of neurons in the second ganglion are larger under strong than weak hypoxia, but both are indistinguishable from conditions of normoxia (p-values from rank-sum tests).The average magnitude of (**F**), number of (**G**), and interval between (**H**) calcium transients of neurons of the second (green symbols and lines) and the third (orange symbols and lines) ganglion in a fish exposed to hypoxic conditions is shown.Light circles are the medians across all neurons in a given ganglion from a given fish; bright squares are averages across fish; bars are the S.E.M; lines are least-square fits.

To characterize these calcium transients, we devised an algorithm to identify the peaks in fluorescence which likely correspond to either bursts or single action potentials (Figure 3D. and see Methods). We found that strongly hypoxic conditions induced transients in the second vagal sensory ganglion that were larger than in weakly hypoxic conditions (p = 0.01); however, the transients under strong and weak hypoxia were indistinguishable from normoxia (Figure 3E). This suggests that under normoxia the transients in this ganglion have highly variable magnitudes, but they tend towards smaller magnitudes under weak hypoxia and larger magnitudes under strong hypoxia. Consistent with this hypothesis, the magnitude of responses in the second ganglion was most strongly correlated with the partial pressure of oxygen (pO_2_) at values of *weight* near 0.7, with a statistically significant (p < 0.05) correlation at these values, and showed larger magnitude validated peaks at lower pO_2_ (Figure 3F, p-value of linear regression is 0.01; Supplementary Figure S4A,B). We also analyzed the average number of transients (Figure 3G) and average length of time between transients (Figure 3H) in both ganglia, but did not find any significant correlation with pO_2_. We also note that normoxia and hypoxia tended to activate separate subpopulations of neurons in either ganglion (data not shown). Collectively, our results indicate that oxygen level modulates the magnitude of calcium events in neurons of the second vagal sensory ganglion in a graded manner.

### Purinergic signaling between NECs and neurons in the second vagal ganglion modifies pectoral fin movements

We next used small molecule agonists to gain insights into the neurotransmitter pathways that may relay oxygen information from NECs to vagal sensory neurons. Previous studies have shown that ATP induces burst activity in the zebrafish petrosal sensory neurons and drives hypoxia-induced respiratory responses [25, 26]. Moreover, zebrafish larvae express purinergic receptors in their cranial sensory neurons and increase their rate of pectoral fin movements in response to ATP agonist [22, 27]. Therefore, we hypothesized that purinergic signaling can activate the vagal sensory neurons and drive pectoral fin movements in larval zebrafish. Animals treated with a P2 purinoceptor agonist, α,β-methyleneadenosine, at either of two concentrations showed an increase in pectoral fin movements (Figure 4A,4B), suggesting that purinergic signaling modifies respiratory behavior. In contrast the agonist increased heart activity only at higher concentrations (Figure 4C, 4D). Next, we tested whether this agonist-evoked behavioral response required neurons in the second or third vagal ganglia. We found that ablating the second, but not the third, vagal sensory ganglion attenuated the agonist-induced increase in pectoral fin movements, suggesting that the second vagal sensory ganglion is activated by purinergic signaling to drive respiratory behavior (Figure 4E, Supplementary Figure S5). On the other hand, ablating neither the second nor third ganglion seemed to have an effect on purinergic modulation of the heart activity (Figure 4F). Taken together, our experiments suggest that the neuroendocrine cell-vagal sensory neuron synapses use purinergic signaling to modulate hypoxia-triggered respiratory behaviors.

**Figure 4.**
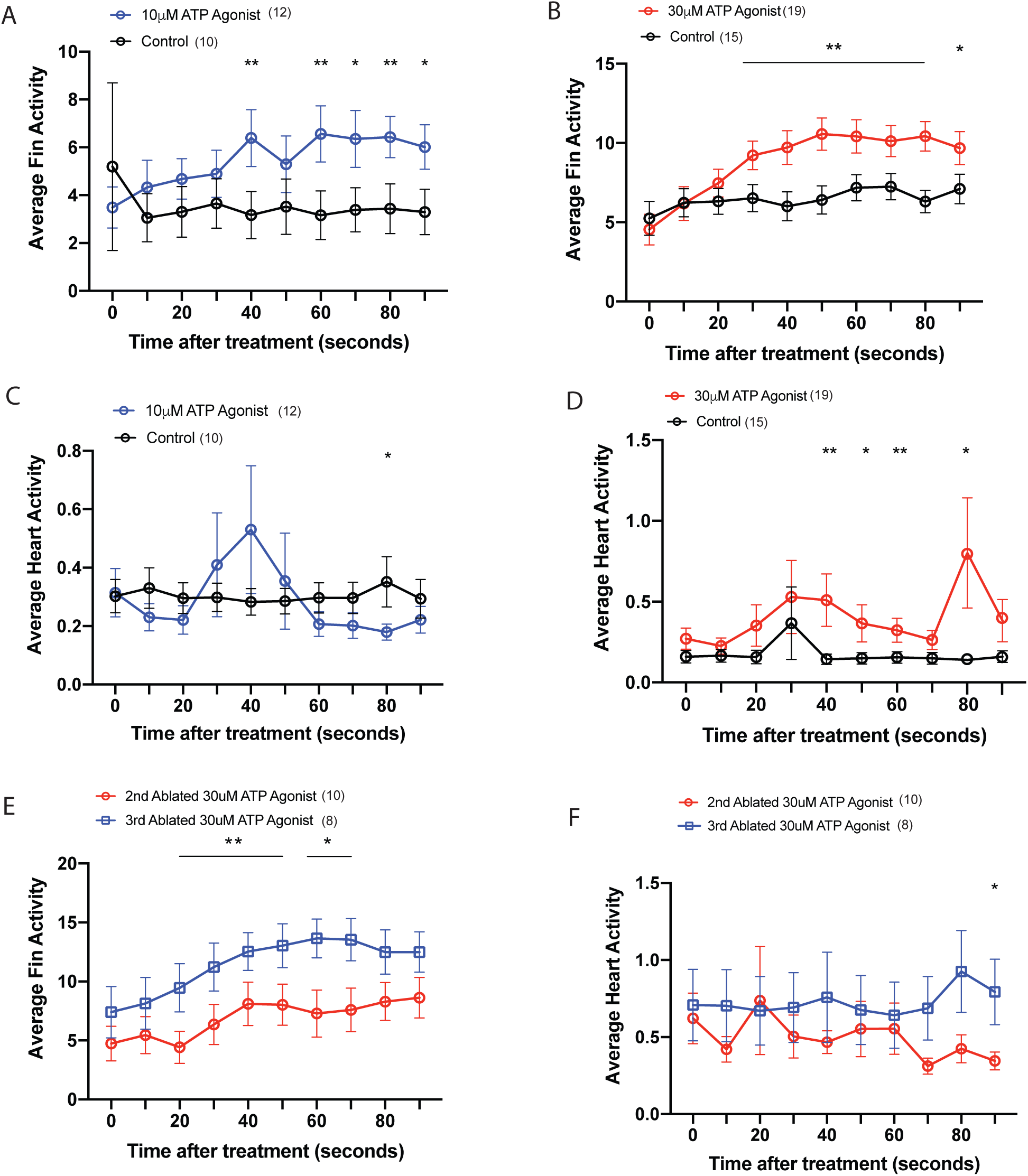
ATP Agonist affect average pectoral fin and heart activity. Average pectoral fin activity in animals exposed to a medium with **(A)** 10 μm or **(B) 30** μm α,β-methyladenossine, P2 purinoceptor agonist. Average heart activity in animals exposed to **(C)** 10 μm or **(D) 30** μm α,β-methyladenossine. Ablating neurons in the second, but not third vagal sensory ganglion blocks the P2 purinoceptor agonist induced increase in (E) pectoral fin, not (F) heart activity. Averages and s.e.m. are shown, n ≥ 8 with * indicating p < 0.1 and ** p< 0.05 using one sample t-test.

## Discussion

Our study shows that vagal sensory neurons respond to changes in environmental oxygen and likely drive hypoxia-induced behaviors in zebrafish larvae. We also suggest that individual vagal sensory neurons receive inputs from multiple NECs and increase pectoral fin movements to facilitate respiration upon exposure to hypoxia. Finally, we find that the magnitude of calcium events in vagal sensory neurons is modulated by pO2. As others have shown that the magnitude of change in GCaMP fluorescence is positively correlated with the level of neuronal activation [28], this suggests that neurons in the second vagal ganglion are more likely to fire more action potentials upon exposure to severe hypoxia, and this may drive an increase in pectoral fin movements.

Where do the vagal axons project? The central projections of the glossopharyngeal and the four vagal sub-ganglia enter the hindbrain at the presumptive rhombomere 6 via a series of nerve roots to form a plexus that could also contains axons of the facial neurons [22]. Interestingly, the posterior half of the vagal motor nucleus is directly dorsal to the pectoral and occipital motor neurons [29]. While the circuitry between vagal axons and pectoral motor neurons has not been mapped, we propose that this connection might play a crucial role in altering pectoral fin movements upon hypoxic activation of vagal sensory ganglia.

We posit that the NEC-mediated purinergic activation of vagal sensory neurons is key to initiating hypoxia-induced respiratory behavior, making it similar to the initiation of nociception, pain, and other mechanical sensations [30, 31]. Neuroendocrine cells in the zebrafish larvae are similar to the Type I glomus cells of the mammalian carotid body. Cell culture and mouse knock-out studies have been used to confirm a central role for ATP and its receptors P2×2/X3 in mediating signaling between the Type I glomus cells and nearby petrosal neurons [32-34]. We showed that ATP agonists increase respiratory behaviors (i.e., pectoral fin movements) in zebrafish larvae, and that this response was blocked by ablating neurons in the second, but not third, vagal sensory ganglia. These results imply that zebrafish NECs also release ATP to activate neurons in the second vagal ganglion. While we are unable to directly monitor activity in the NECs, we speculate that hypoxia depolarizes these cells, which then leads to a calcium-dependent release of ATP and subsequent activation of the downstream neurons in the second vagal ganglion. Future studies should aim to test this proposed circuit.

We monitored changes in pectoral fin movements in animals experiencing hypoxia. Pectoral fins are not required for the initiation of swimming or for rhythmic swimming behaviors in zebrafish larvae. Rather pectoral fin movements facilitate gas exchange near the skin of the animal [12]. Our results are consistent with a role for pectoral fins in respiratory behavior. We suggest that larvae exposed to hypoxia in our microfluidic devices increase their pectoral fin movements to bring water with higher pO_2_ caudally over the fins to increase their exposure to oxygen. Furthermore, when these efforts are unsuccessful, they increase their tail and posterior body movements to escape (“delayed startle responses” [35]). In summary, our analyses suggest a role for purinergic signaling at the many oxygen-sensing neuroendocrine cells-vagal sensory neuron synapse to drive respiratory behaviors in response to hypoxia. We also suggest that further analysis of this circuit will reveal additional insights into signaling pathways that may be conserved across species.

## Methods

### Calcium imaging and analysis

The imaging setup was described previously [36]. Briefly, single 5 days post fertilization (d.p.f.) *elavl3*:*H2B-GCaMP6f* [24] larva with the opercula removed 1 day prior was paralyzed with a drop of 1mg/ml alpha-bungarotoxin and then embedded in the capillary with 1% low-melting agarose. A peristaltic pump was used to exchange the media around the larva, switching from normoxic to hypoxic media (Figure 3A). We used the light-sheet microscope system to excite neurons within the cranial ganglia and captured the resulting emission at 90ms/frame (Figure 3B). pO_2_ levels were measured in real time and datasets where these levels deviated more than 10% from the expected value were excluded. Changes in fluorescence from individual neurons were obtained and denoised after correcting for motion using the MATLAB-based tool CalmAn [37-41]. MATLAB functions **findpeaks** was used to detect fluorescence transients. We detected peaks by looking for local maxima that had a prominence at least as large as 1) *weight* multiplied with the standard deviation of the time series and 2) *weight* multiplied with the mean of the time series. When the tunable parameter *weight* is set low, relatively weak peaks are counted as spikes; this may incorrectly classify noise as a spike. On the other hand, when *weight* is set high, relatively prominent peaks, which may have resulted from actual spikes, may be rejected (Supplementary Figure S3). Hence, we performed a sensitivity analysis to see how our results depend on the value of *weight* (Supplementary Figure S4). Specifically, we varied *weight* from 0.5 to 0.8 with 0.01 increments and assessed the strength of responses to various levels of hypoxia in terms of the average magnitude of detected peaks. We excluded (1) all recordings where less than 2 neurons were active and (2) all neurons with fewer than 2 transients during the recording window. The alpha value for testing the significance of each regression was adjusted by the effective number of independent variables, or *m*_*eff*_, which was computed using a matrix spectral decomposition approach to account for the fact that two of our variables of interest were likely correlated (i.e., the number of peaks and the interpeak interval [42]) Hence, alpha was 0.02 (= 0.05/*m*_*eff*_, where *m*_*eff*_ = 2.5). It is important to note that, even though the linear regression was not statistically significant at lower and higher values of *weight*, the negative correlation between pO_2_ and magnitude of response in the second ganglion always remained, suggesting a robust trend (Supplementary Figure S4C). It is also worth noting that there was never a significant correlation between pO_2_ and magnitude of calcium peaks in the third ganglion at any value of *weight* (Supplementary Figure S4A,B), nor was there a significant correlation in the second ganglion under normoxia (Supplementary Figure S4E), suggesting that a simple tuning of *weight* does not lead to a spuriously significant result.

### Immunohistochemistry

Larvae were fixed in 4% paraformaldehyde with regions of interest labeled using the following primary antibodies: goat anti-GFP (1:200; Abcam), rabbit anti-serotonin (1:200; ImmunoStar), rat anti-serotonin (1:200; Novus Bio), or rabbit anti-Kaede, (1:200; MBL), which in turn were detected using secondary donkey anti-goat Alexa Fluor 488 (1:1000; Invitrogen), goat anti-rabbit Alexa Fluor 546 (1:1000; Invitrogen), or donkey anti-rat Alexa Fluor 546 (1:1000; Invitrogen). Larvae between 4-6 d.p.f. were imaged in 70% glycerol using a Zeiss LSM800 or Olympus FM1000-MPE system. Subsequent image processing and evaluation were performed in ImageJ.

### Microfluidic device

The microfluidic device for imaging zebrafish larvae under an inverted microscope is assembled out of three parts of laser-cut acrylic and a cover glass (Figure 1A). The thicknesses of acrylic in parts 1, 2, and 3 are 0.75, 0.75, and 1.5 mm, respectively. To bond the parts together, part 1 has an ∼0.13 mm layer of pressure sensitive adhesive (PSA) at the bottom, and part 2 has layers of the same PSA on both sides. After part 1 is bonded to the cover glass, an ∼5 mm diameter, ∼0.9 mm deep round cuvette is formed, where a Tricaine-anesthetized zebrafish larva is trapped inside 2.5% low-melting agarose gel (UltraPure LMP Agarose, Invitrogen 16520-050). To facilitate the exposure of NECs to oxygen in the medium and observe larval movements, the agarose gel is removed from areas around the gills, pectoral fins, and tail using a thin needle. After parts 2 and 3 are bonded to the top of part 1, a sealed perfusion channel with two inlets and one outlet (holes in part 3) is formed directly above the trapped animal. The inlets and outlets are connected to separate medium reservoirs through lines of flexible PVC tubing. The tubing is compression inserted into laser-cut acrylic ports (∼3 mm thick, ∼6 mm in diameter) that have sticky rings of ∼0.6 mm thick PSA at the bottom, and the ports are bonded to part 3, placing the tubing lines coaxially with the inlets and outlet (Figure 3A). The flow through the device is driven by a peristaltic pump connected to the outlet. At any given time, only one of the inlet tubing lines is open, while the other is blocked (Figure 3A).

The microfluidic device for imaging of zebrafish larvae under a light sheet microscope is assembled out of four parts of laser-cut acrylic and has two small thin windows, for the imaging objective and light sheet (Figure 3A). The thicknesses of acrylic in parts 1 – 4 are 0.2, 1.5, 0.5, and 0.25 mm respectively. The imaging window is 0.05 mm thick and made of COP, and the acrylic light sheet window is 0.2 mm thick. To bond the parts together, part 2 has an ∼0.13 mm layer of a PSA at the top, and part 3 has layers of the same PSA on both sides. After the imaging window is glued to part 1, and parts 1 – 3 are bonded together, the light sheet window is glued to the front, and the entire assembly is turned upside down. A small amount of 2.5% low-melting agarose gel with *Tricaine* is dispensed into the cuvette formed by the cavities in parts 1 and 2, a *Tricaine*-anesthetized zebrafish larva is placed into the cuvette, and the excess of gel pre-polymer is removed, leaving the larva barely covered. Solidified gel is removed from areas around the gills, pectoral fins, and tail using a thin needle. Part 4 is bonded to part 3, making a sealed microfluidic device with two inlets and one outlet and with a perfusion chamber (part 3) beneath the partially immobilized larva. The inlet and outlet ports with PVC tubing (same as in the device in the previous paragraph) are bonded to the top (Figure 3A).

### Behavioral analysis

All the behavioral data were recorded from 5 d.p.f. wildtype *wik* strain or *Tg(p2rx3*.*2:gfp)*^*sl1*^ strain [22] raised under 14/10 light cycles in E2 medium. An oxygen meter (PreSens: Fibox 4, sensor: Cat. 200000241 and cell: Cat. 200001690) was attached to the outlet to measure the oxygen concentrations of the medium in real time (5-10 sec/read). The recording setup consists of a Basler aca1300 camera (60 fps, Edmond Optics), illuminating base (Tritech Research, stereo-microscope base) with a set of lenses (Scheider Inc., catalogue #s 25-1070160, 25-020178, 25-011726, 25-020155, and 25-020052) and Noldus EthoVision XT10 software. Individual regions of interest were assigned to the pectoral fins and heart (Figure 1C) and the movements within the regions were detected by Noldus software (heartbeat detector). The percentage of the pixels that change intensity in individual regions were defined as activity. Pectoral fin and heart activity were analyzed. Hypoxia was induced by feeding into the microfluidic device a medium equilibrated with a hypoxic gas mixture (E2 bubbled with nitrogen/air mixture which reduces pO_2_). The pO_2_ range was 130 mmHg for normoxia, 60-80 mmHg for mild hypoxia and 10-35 mmHg for strong hypoxia. Osmotic shock was induced by adding 100 mM or 250 mM sodium chloride solution into the microfluidic device. For drug treatments, α,β-methyleneadenosine 5’-triphosphate trisodium salt (TOCRIS: Cat. 3209), hydrochloride were dissolved into 100 mM stock in Milli-Q water and further diluted into different concentrations. Each larva was trapped and its behavior monitored with control solution before being replaced with drug-containing solutions.

### Laser ablation

Ablations were performed on *Tg(p2rx3*.*2:gfp)*^*sl1*^ larvae at 3 d.p.f. Larvae were anesthetized with Tricaine and mounted laterally on cover slides. Tricaine was used at a working dilution of 1.2ml in 20ml of E2 medium (Stock 4g/1000 ml, Tricaine MP Biomedicals #103106). Imaging and ablation were performed with Olympus FM1000-MPE system with 25x water immersion lens (XLPL25XWMP). An 800 nm laser beam was focused onto individual cranial sensory ganglion for 5 seconds, and clear lesions were detected. The ablation of neurons was confirmed before behavioral experiments.

### Statistics

Unless stated otherwise, data points represent averages of all data and error bars are standard error of the mean (S.E.M.). Two-tailed one sample *t-*test were used to test the differences between the control and treated groups. Two-tailed Student’s *t*-tests or Mann-Whitney *U*-tests (when the data did not comply with the assumptions of the *t*-test) were used for two-group comparisons. ANOVAs were used for multiple group comparisons, followed by Bonferroni’s or Sidak’s multiple comparison test. Prism 8 (GraphPad) was used for other analysis and generating plots. Unless stated otherwise, ^*^ indicates *p* < 0.1, ^**^ indicates *p* < 0.05.

## Supporting information

Legends for Supplementary Videos

## Data and Code Availability

The code used to analyze calcium imaging data along with the data are available at https://github.com/shreklab/Rosales-Yeh2021. Further information and requests for resources and reagents should be directed to and will be fulfilled by the Lead Contact, Sreekanth H. Chalasani.

## Supplementary Data

Supplementary Information includes five supplementary figures (S1-S5), and four supplementary videos (S1-S4).

## Acknowledgements

We thank D. Thai and a number of UCSD undergraduates who helped with zebrafish care and maintenance. We also thank L.A. Hale and M. Erickstad for inputs into our behavioral setup, Uri Manor and the Waitt Advanced Biophotonics Center for imaging resources, W. van Giessen for technical advice on analyzing behavior data, C. Fernandes for illustrations, H. Burgess, S. Navlakha, J. Fetcho, G. Haddad, S. Ryu, E. Mukamel, J. Wang, D. O’Keefe, C. Lee-Kubli, M. Matty, M. Rieger, and A. Singh for comments on the manuscript. We are grateful to all the members of the Chalasani laboratory for critical help, advice and insights. C-M. Yeh was supported by a Salk Pioneer Fellowship. This work was funded by innovation grants from the Kavli Institute of Brain and Mind, UCSD (C-M.Y and G.M.P), and the Salk Institute for Biological Studies (S.H.C.), NSF award 1411313 (A.G.). The Waitt Advanced Biophotonics Center is supported by NCI CCSG P30014195.

## Author Contributions

K.R. performed experiments and analyzed data. C-M.Y. conceived, designed, and performed experiments. J.J.H. analyzed the data. R.V-R., and E.D performed experiments, A.G. designed the microfluidic device and co-wrote the manuscript, and S.H.C. conceived the experiments and co-wrote the manuscript.

## Declaration of interests

The authors declare no competing interests.

## Figure Legends

Legends are included in the figures.

## Notes

### Competing Interest Statement

The authors have declared no competing interest.

### Summary of Updates

Section on neural coding is updated. Figures 1-4 revised. Authors and Supplemental files updated

