## Supplementary material for "Neural pathways linking hypoxia with pectoral fin movements in *Danio rerio*": Legends for Supplementary Videos

Supplementary Information includes five supplementary figures (S1-S5) and four supplementary videos (S1-S4).

Figure legends for Videos are included in this file, while those for each of the Supplementary figures are at the bottom of each figure.

**Supplementary video S1**

Example video showing behavioral responses of a 5 dpf larva to normoxia stimulus.

**Supplementary video S2**

Example video showing behavioral responses of a 5 dpf larva to 60-80 mmHg hypoxia stimulus.

**Supplementary video S3**

Example video showing changes in GCaMP6f fluorescence from neurons in the 2$^{nd}$ and 3$^{rd}$ vagal sensory neurons of a 5 dpf larva under normoxia.

**Supplementary video S4**

Example video showing changes in GCaMP6f fluorescence from neurons in the 2$^{nd}$ and 3$^{rd}$ vagal sensory neurons of a 5 dpf larvae under 50 mmHg hypoxia.
