## Supplementary Figure 1 for "Neural pathways linking hypoxia with pectoral fin movements in *Danio rerio*"

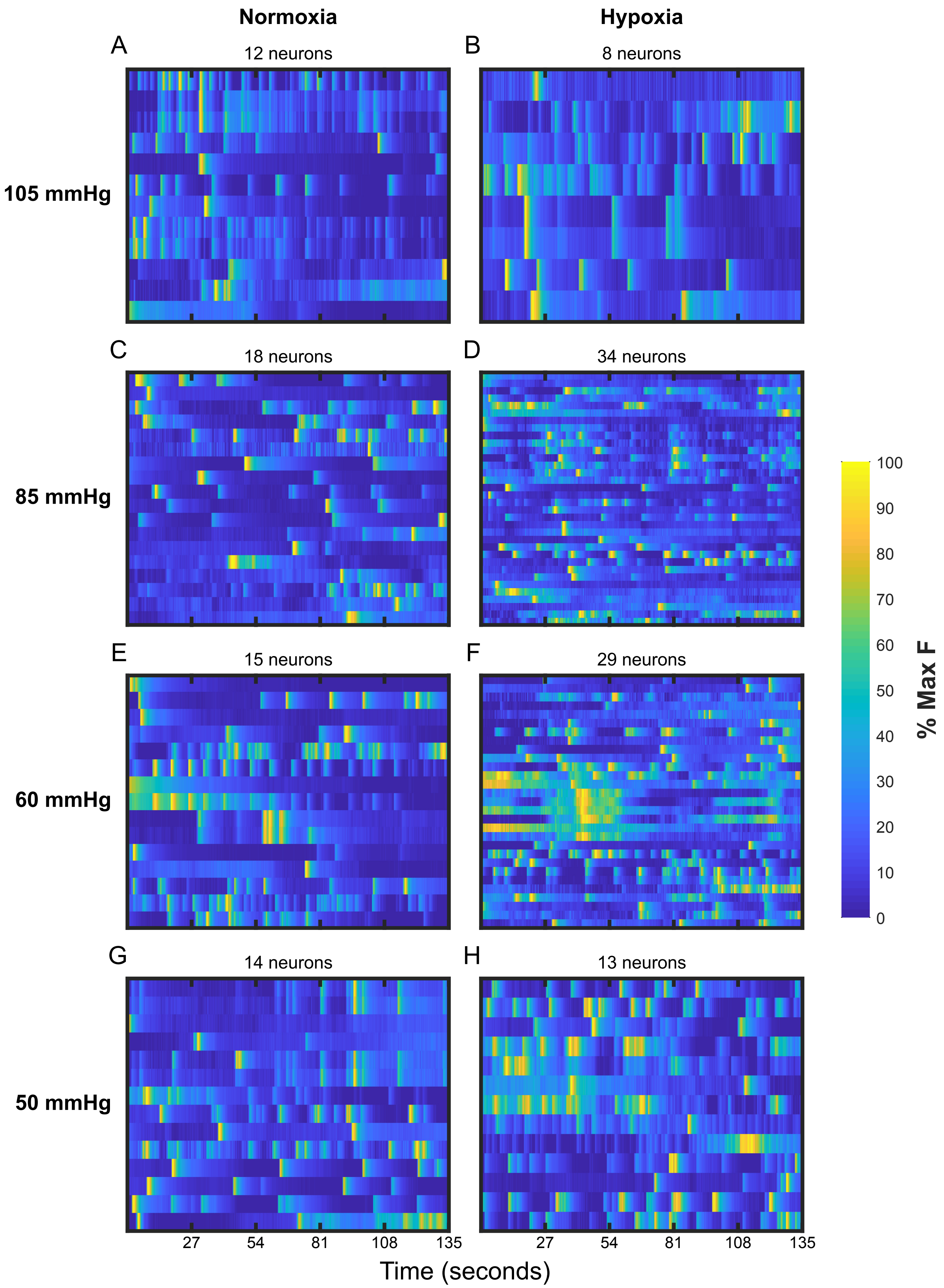


**Supplementary Figure 1 | Fluorescence of neurons in the second vagal ganglion of several fish.** The responses of recorded neurons to normoxia (**A**, **C**, **E**, **G**) and hypoxia (**B**, **D**, **F**, **H**) are shown for different partial pressures of oxygen (i.e. 105, 85, 60 and 50 mmHg). Partial pressure of oxygen is always the same for normoxia sessions, which serve as controls, and only varies for the hypoxia sessions. Responses are normalized by the cell's maximum fluorescence, and the number of cells analyzed are shown above each heat map. Different numbers of neurons were active in normoxia and various partial pressures of hypoxia.
