## Supplementary Figure 3 for "Neural pathways linking hypoxia with pectoral fin movements in *Danio rerio*"

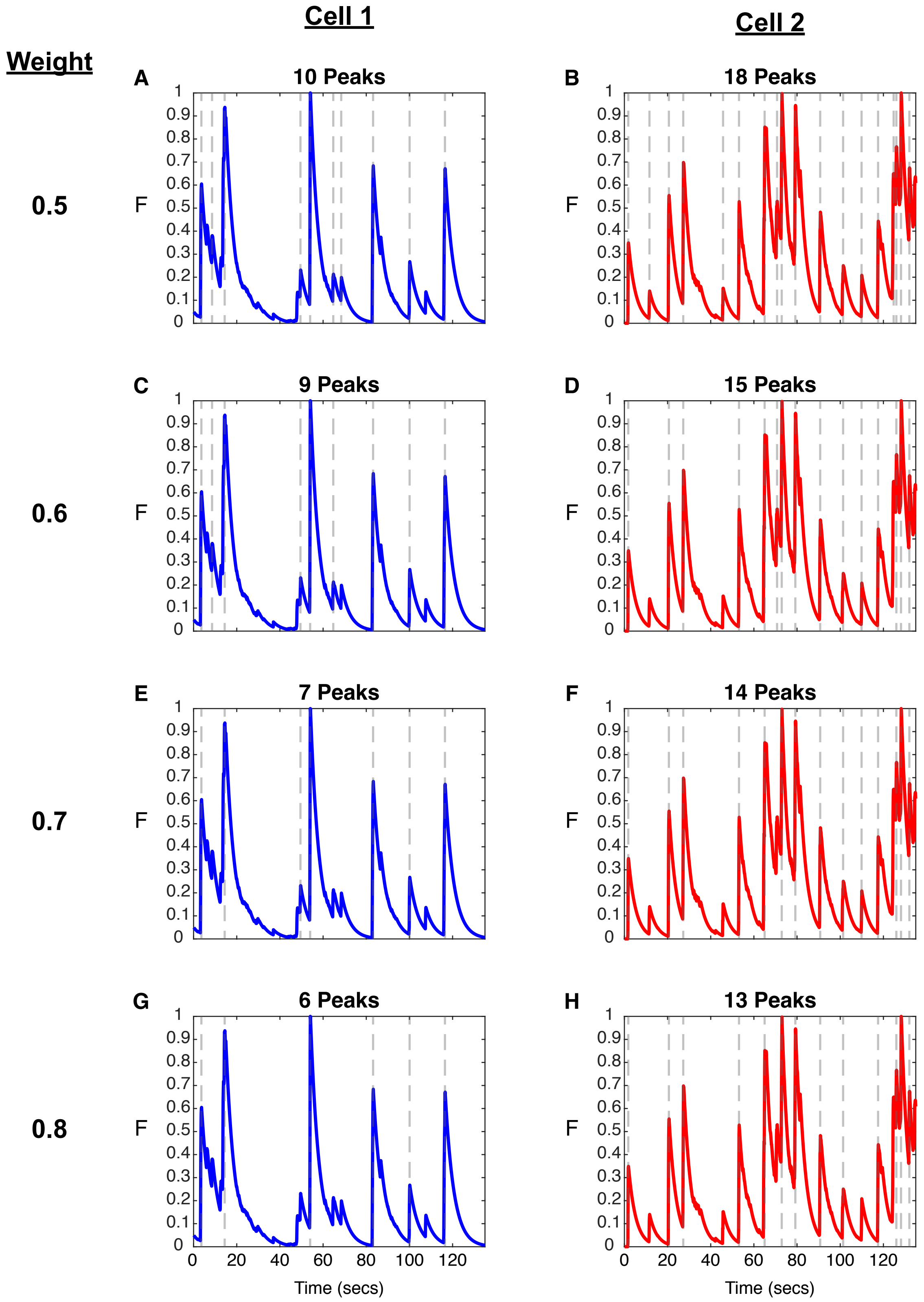


**Supplementary Figure 3 | The detection of calcium transient is sensitive to the weight parameter.** Traces from two cells (i.e. cell 1 in **A**, **C**, **E** and **G** in blue, and cell 2 in **B**, **D**, **F**, and **H** in red) have several peaks (indicated by dotted grey lines). There are more peaks at lower weight (e.g. **A** and **B** vs **G** and **H**) because the less stringent requirements include lower magnitude peaks, potential noise, or both. Traces are normalized to their maximum fluorescence value.
