## Supplementary Figure 4 for "Neural pathways linking hypoxia with pectoral fin movements in *Danio rerio*"

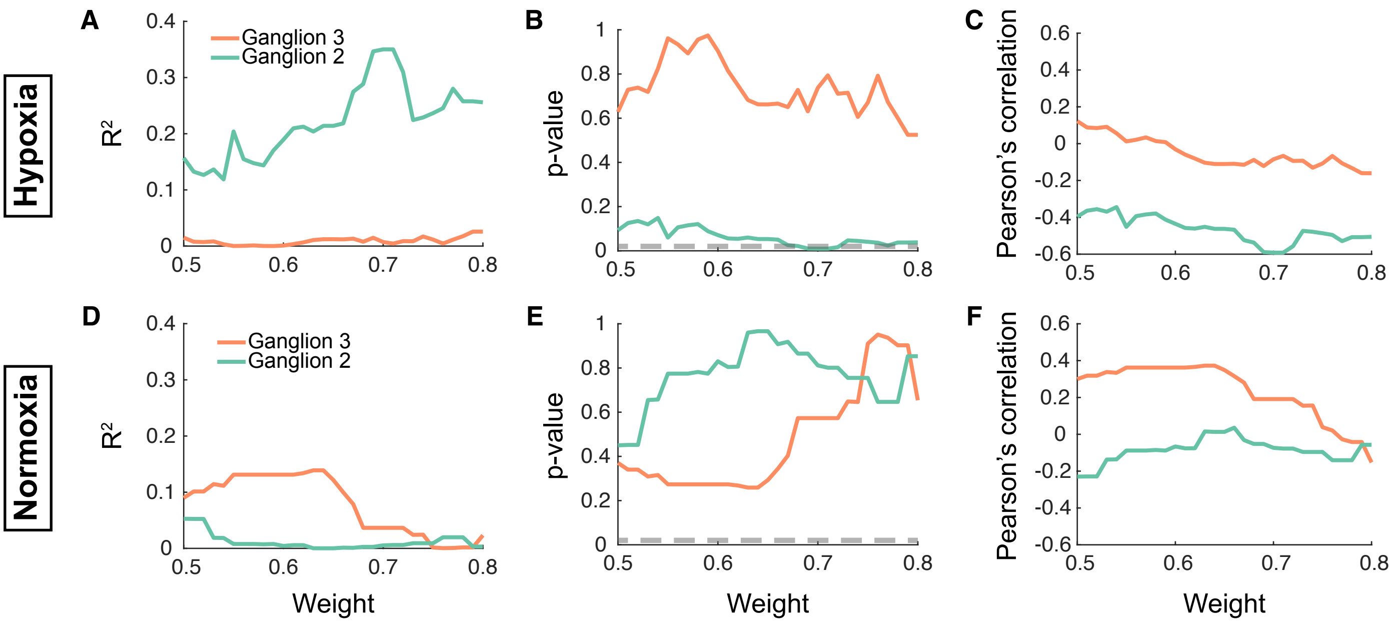


**Supplementary Figure 4 | Sensitivity analysis shows that hypoxia affects the magnitude of high-confidence calcium transients in the 2nd ganglion.** Every panel describes the linear relationship between magnitude of calcium transients and four levels of hypoxia in the 2nd (green) or 3rd (orange) ganglion as a function of the weight used in our algorithm to find transients; the higher the weight, the less likely any detected transients, or peaks, were noise. Panels (**A** and **D**) depict the R2 of this linear regression, while panels (**B** and **E**) show the corresponding p-value, at different weights. Panels (**C** and **F**) show the Pearson's correlation coefficient between magnitude and level of hypoxia at different weights. Animals experienced a hypoxia and normoxia session, and were assigned to just one level of hypoxia. Panels (**A** – **C**) are for hypoxia sessions, while panels (**D** – **F**) are for normoxia sessions. The dotted grey line in panels (**B** and **E**) correspond to the adjusted alpha of 0.02 (= $0.05/m_{eff}$ , where $m_{eff}$ = 2.5); p-values below this line indicate a statistically significant linear relationship between magnitude and levels of hypoxia.
