## Supplementary Figure 5 for "Neural pathways linking hypoxia with pectoral fin movements in *Danio rerio*"

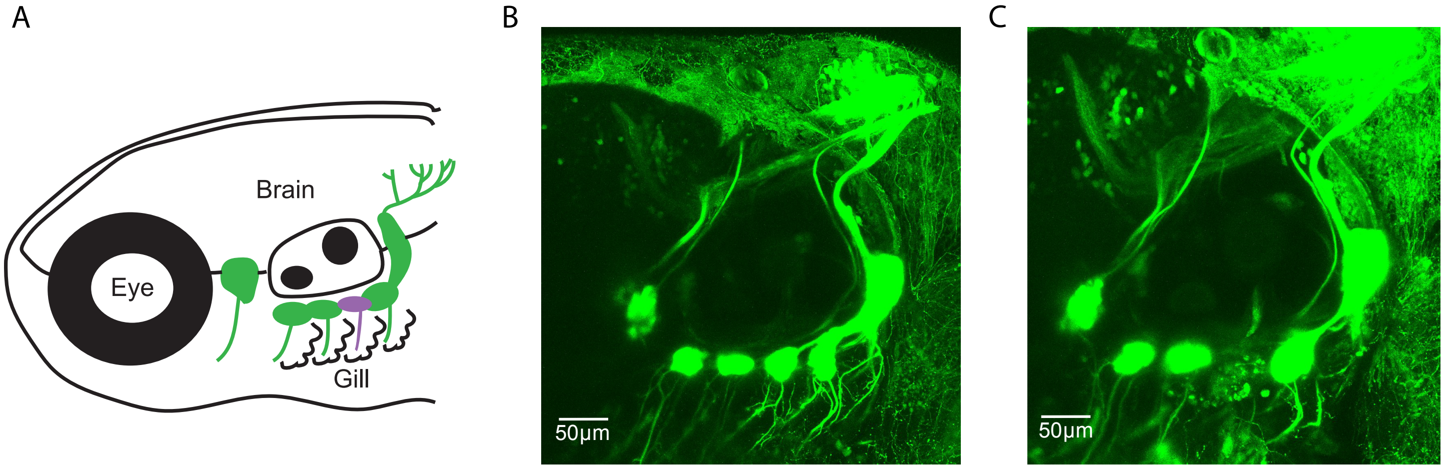


**Supplementary Figure 5 | Ablations to delete neurons in vagal sensory ganglion.** (**A**) Schematic showing the position and orientation of the larvae and relevant structure. Images showing a lateral view of a *Tg(p2rx3.2gfp)* fish (**B**) before and (**C**) after neurons in the second vagal ganglion are ablated.
